# Driving factors for beta-lactam resistance gene amplification during *de novo* resistance evolution in *E. coli*

**DOI:** 10.1101/2025.02.07.637019

**Authors:** Luyuan Nong, Xinwei Liu, Xinyu Wang, Wim de Leeuw, Martijs Jonker, Stanley Brul, Benno ter Kuile

## Abstract

Long-term exposure of *E. coli* to non-lethal step-wise increasing concentrations of beta-lactam antibiotics induces high levels of resistance that can be accompanied by amplification of a chromosomal fragment around the *ampC* gene. We compared the amplification of the *ampC* fragment in the wildtype, an *ampC* knockout mutant, a mutant in which the *ampC* gene was replaced by a tetracycline resistance gene *tet(B)*and a strain in which the *ampC* has been translocated. When *ampC* was removed, no amplification occurred at the original *ampC* location, but DNA fragments were amplified around the genes coding for efflux pump AcrAB and the multiple antibiotic resistance operon MarRAB. When *tet(B)* replaced *ampC*, exposure to tetracycline induced amplification of comparable fragments, while exposure to amoxicillin induced duplication of a larger fragment elsewhere. When *ampC* was translocated, a fragment around it at the new location was amplified. The importance of the presence but not of the location within the chromosome of the resistance genes for the amplification process indicates that the mechanisms are neither gene nor location specific. Without the relatively efficient resistance gene *ampC*, duplication and amplification occur around *acrAB* and *marRAB* that code for amoxicillin and tetracycline resistance factors. These duplications and amplifications are prevented by *ampC* amplification.

## Introduction

Antimicrobial resistance (AMR) in microbes causes considerable risks for human health care (1). Resistance development can be caused by adaptation at the cellular level, DNA mutations due to long-term exposure to sublethal levels of antimicrobials and horizontal transfer of resistance genes. The consequences for human health care of the exchange of resistance plasmids between bacteria making pathogens resistant have been documented extensively (2). The *de novo* mechanism for building up antibiotic resistance at the genetic level may over time interact with the plasmid-bound systems when resistance genes are amplified and end up in fragments that can subsequently be transferred to previously susceptible cells (3, 4). Horizontal gene transfer is generally considered to be the main cause of antimicrobial resistance spreading and prevalence.

When bacteria are exposed to sublethal levels of antibiotics, the cell’s regulatory network is activated to adapt the microbe to this environmental change (5), generating *de novo* resistance in naive wild-type strains (6). In the case of *E. coli* exposed to amoxicillin, after initial adaptation (7), a mutation selection process of trial and error at the single nucleotide level (8) and genetic rearrangements (9) occur. These genetic rearrangements include the amplification of a fragment around the *ampC* gene, resulting in a flanked-by-transposon fragment that can be transferred to an acceptor susceptible cell, that becomes resistant as a result (4).

Duplication events have a higher frequency than point mutations, but the reversion rates are so high that the outcome is a steady state (10, 11). The overall effect is that resistance gene duplication and amplification (GDA) in an evolving populations is sometimes observed only after point mutations accumulation (12, 13). The dynamic interaction between GDAs and point mutations causes the preservation of some of the amplification events, forming the basis of evolutionary novelties (14).

The GDAs and point mutations accumulated are conserved in some species during long-term resistance evolution (3, 15, 16), especially if there is a moderately efficient resistance gene in the chromosome. During *de novo* evolution of amoxicillin resistance in *E. coli,* the beta-lactamase AmpC activity and the MIC exhibited a linear relationship (17). The *ampC* gene has low expression levels as it is regulated by a weak promoter in wild-type *E. coli*. After resistance evolution, the prevalent GDAs and multiple point mutations are related to the *ampC* gene, resulting in a more than 100-fold increase in *ampC* transcription (3, 17). The size of these amplicons varies between 2.6 and 13.4 kb, and their boundary sites are located at a limited number of locations (4, 9, 16–18). Hence, it is possible that the location of *ampC* gene impacts the GDA event. When an *ampC* knockout mutant (Δ*ampC*) was subjected to the same experimental procedure, the GDAs and point mutations observed in the WT did not occur. Moreover, compared to the WT, several point mutations were either absent or increased in frequency during the evolution of Δ*ampC* (17).

The GDAs can arise by homology or non-equal homology recombination between sister chromosomes via RecA-dependent or RecA-independent mechanisms (19). In some cases, GDAs can also be mediated by RNA reverse transcription (20) and transposon elements. However, there is controversy about how the transposon elements facilitate GDA events, depending on homology recombination (21, 22) or transposition (23, 24). The junction between the tandem connection of GDAs can be used to analyse its putative mechanism(s).

The mechanisms of antibiotic resistance in bacteria include among others antibiotic inactivation, decreased influx, active efflux, target protection, and many more (25). Each pathway carries different fitness cost. The modification or inactivation of the antimicrobial drug itself, like AmpC, is associated with less fitness costs than other mechanisms, such as the ATP-binding cassette (ABC) transporters, which facilitate multidrug resistance (26). When fitness costs of a resistance mechanism are high, cells have a selective advantage if the changes that cause resistance can be reversed when the antibiotic is no longer present.

When *E. coli* is exposed to step-wise increasing concentrations of a beta-lactam antibiotic, a fragment of its chromosome around the *ampC* gene is amplified (4). This fragment has a limited number of starting and ending points and can be transferred to a naive wildtype cell that upon incorporating it, becomes resistant as well. What is not clear is whether the cellular mechanisms that achieve the GDAs in *de novo* beta-lactam resistance development are in some way “aware” of the usual location of *ampC* in the chromosome, or that the amplification is triggered by increased rates of transcription. In the first case removing or translocating the *ampC* gene would not affect the GDA of the fragment, and it would still proceed, even though it lacks *ampC*. If the second mechanism, for which there is experimental evidence (12, 14), is closer to reality, replacing the *ampC* gene with a relatively weak resistance gene for another antibiotic, would upon exposure to that antibiotic still induce GDA.

Here we investigate the change in occurrence of GDAs and point mutations during resistance evolution when the *ampC* was either deleted, translocated within the chromosome or replaced by the tetracycline resistance gene *tet(B)*. The following questions are addressed:

1. Does the location of a resistance gene influence the incidence and size of the GDAs?
2. Do the GDAs still occur if *ampC* is replaced by a resistance gene for another antibiotic with higher fitness costs?
3. How do the accompanying point mutations change in evolved strains with a different or without a resistance conferring gene?

## Results

### Evolution towards amoxicillin and tetracycline resistance

In order to answer the questions asked in the introduction, three mutants were created: an *ampC* knockout mutant (Δ*ampC*), a strain in which the *ampC* gene was replaced by a tetracycline resistance gene *tet(B)* (TCR), and a translocated *ampC* complementation strain (CompA). These mutants were exposed to step-wise increasing concentrations of amoxicillin or tetracycline to induce resistance (Fig.1a, Fig.2a). The wild type (WT) was used as a biological control to compare the rate of resistance development and cross resistance. For each strain a zero-exposure control was passaged in drug free medium in an otherwise identical procedure.

**Fig. 1.**
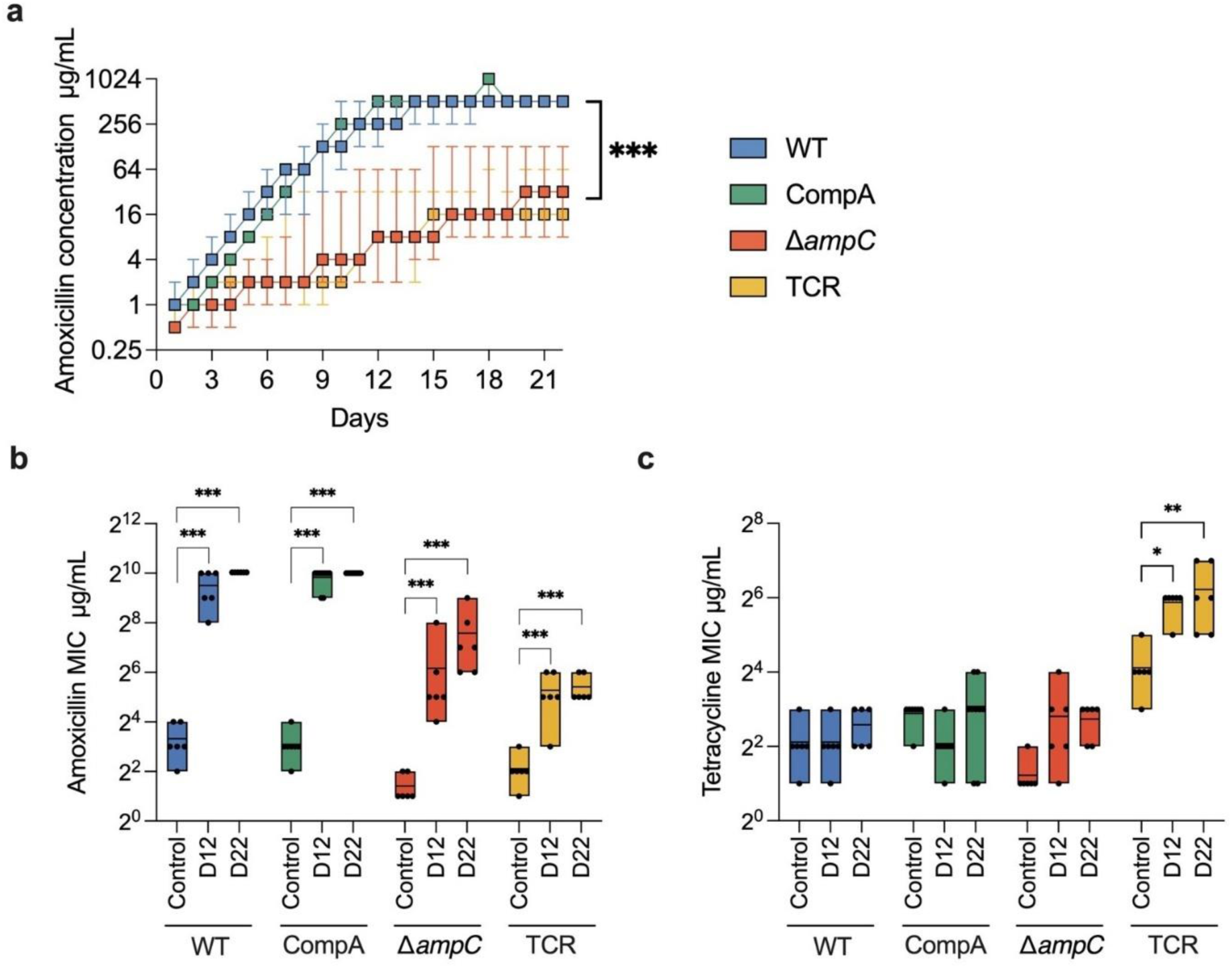
Amoxicillin resistance evolution. **a** The concentration of amoxicillin that still allows growth during stepwise-increase resistance evolution. **b** The MIC of amoxicillin of the control, the evolved strains on day 12 (D12) and day 22 (D22) during the evolution. **c** The change of tetracycline MIC. Different strains are shown in different colors. The Wilcoxon signed-rank test was used for statistical significance analysis of the differences between rates of adaptation. The ANOVA was used for the log_2_ transformed MIC. The significance of the differences between groups of the same strain is indicated with asterixis: \*\*\**p*<0.001, ** *p*<0.01, * *p*<0.05.

**Fig. 2.**
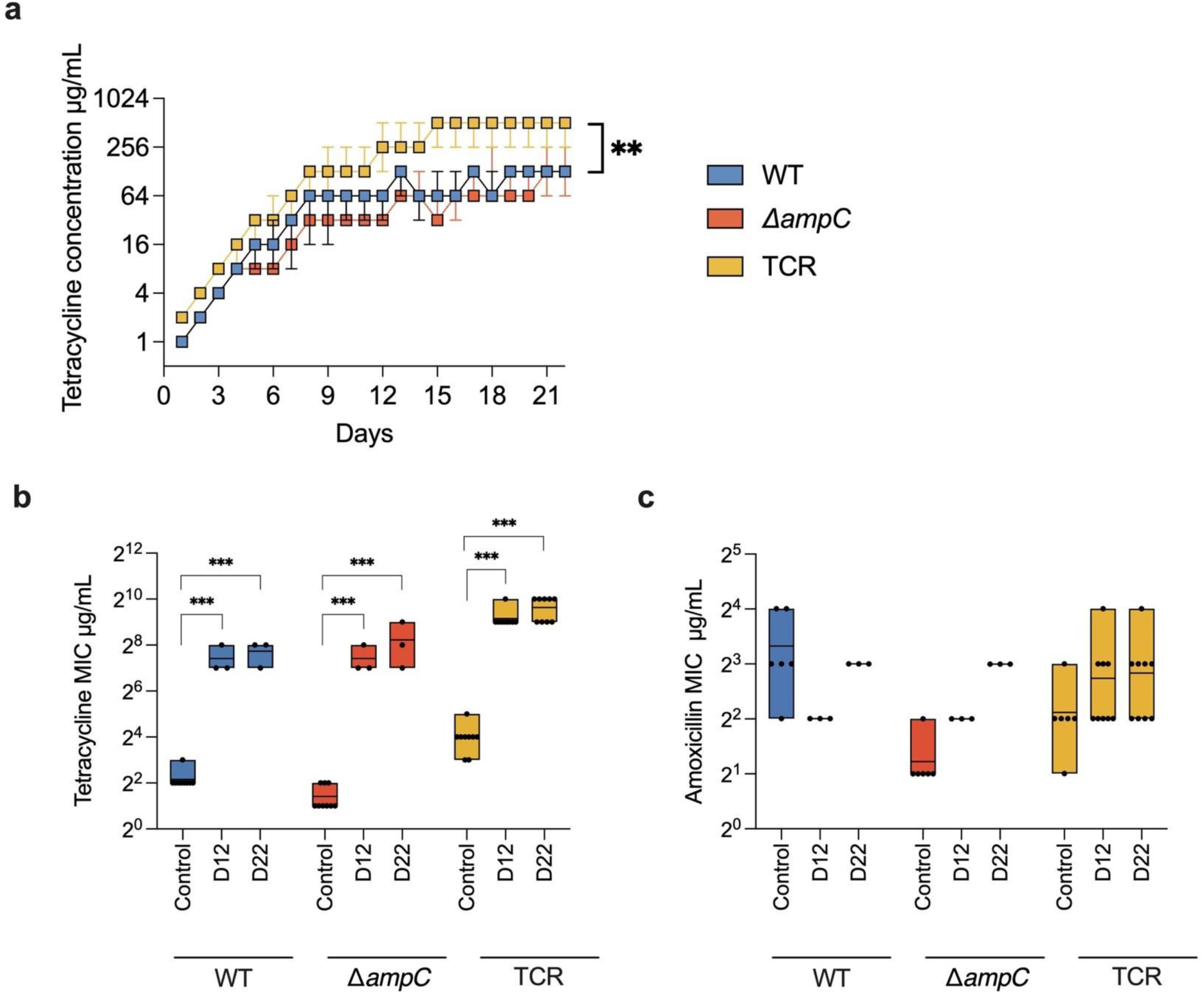
Tetracycline resistance evolution. **a** The tetracycline concentration that still allows growth during the resistance evolution experiments. **b.** The MIC of tetracycline of the unexposed naive strains, the evolving strains on day 12 (D12) and day 22 (D22). **c** The change of amoxicillin MIC. Different strains were shown in different colors. The Wilcoxon signed-rank test was used for statistical significance analysis of adaptive rates. The ANOVA was used for the log_2_ transformed MIC. The significance between groups in the same strain is shown with asterixis. ** *p*<0.01, *** *p*<0.001.

The original WT and CompA both have an 8 mg/L amoxicillin MIC in LB medium. The naive Δ*ampC* mutant has a 4-times lower amoxicillin MIC (2 mg/L). The MIC of TCR is 4 mg/L, which is higher than that of Δ*ampC* but less than that of the WT and CompA, that does possess *ampC*. Like the Δ*ampC,* the TCR strain does not contain the *ampC* gene. Hence the *tet(B),* which codes for an efflux pump, seems to contribute to amoxicillin resistance, but at a lower efficiency than *ampC*.

When exposed to stepwise increasing amoxicillin concentration, the adaptive rates of WT and CompA were similar and high (Fig.1a). All replicates of WT and CompA built up resistance to amoxicillin to the same extent (512 mg/L, Fig.1b). When the *ampC* gene is absent, in the TCR and Δ*ampC* mutants, resistance to amoxicillin was acquired at a much slower rate, but there was still a considerable increase in MIC over the entire experiment. Rather than reaching the same high-level MIC as the *ampC* containing strains, the TCR and Δ*ampC* strains adapted to a wide range of amoxicillin concentrations, varying between 8 to 128 mg/L.

When the TCR strain was exposed to sublethal levels of amoxicillin, the resistance to tetracycline increased (Fig.1c), but not as much as when the cells were exposed to tetracycline (Fig. 2b). In contrast, the resistance to tetracycline of the Δ*ampC* exposed to amoxicillin did not increase. The WT and CompA exposed to amoxicillin did not increase tetracycline resistance either (Fig.1c), thus exposure to amoxicillin only affected tetracycline resistance if the *tet(B)* gene is present. Hence, the increase in tetracycline resistance of amoxicillin resistant TCR is not the result of cross-resistance, but of the overexpression of *tet(B)*.

Exposure to tetracycline induced resistance to that antimicrobial. The naive WT, Δ*ampC*, and TCR had tetracycline MIC’s of 4, 2 and 16 mg/L respectively. The WT and Δ*ampC* adapted at similar rates during tetracycline resistance evolution, while the TCR mutant had a significantly higher rate (Fig.2a). At the end of the evolution experiment, the resistant TCR adapted to higher tetracycline concentrations than WT and Δ*ampC* (Fig.2a). This suggests that the *tet(B)* gene provides a selection advantage under tetracycline exposure. Compared to amoxicillin resistance evolution (Fig.1a), the differences of adaptive rates towards tetracycline resistance of strains with and without the *tet(B)* resistance gene were less. Amoxicillin resistance did not significantly increase during exposure to tetracycline in TCR and WT, but Δ*ampC* developed slightly higher resistance (Fig.2c).

### Fitness of resistant strains

In order to study the fitness costs caused by these different resistance mechanisms, the growth rates of resistant strains were measured with and without antibiotic on day 10 and day 22 of the evolution. Growth rate was used as an indirect measure for fitness. After the resistance evolution, a 15-day reverse evolution, passaging the strains in drug free medium, was conducted. The corresponding MIC was measured every three days.

The development of amoxicillin resistance did not impact the growth rate of WT during the whole evolution process (Fig.3a). Combined with the observation that its reverse evolution did not decrease the resistance (Fig.3c), the *ampC* overexpression seems to carry little fitness cost in the WT. The growth rate of CompA was decreased at day10 at ¼ amoxicillin MIC, but only slightly less than the naive strain at day 22. However, the amoxicillin resistant CompA did not exhibit resistance decrease after reverse evolution. This indicates that the fitness costs of *ampC* overexpression in CompA are also small but higher than those of the WT. On the contrary, the fitness costs of amoxicillin resistance in Δ*ampC* and TCR were higher than those in WT and CompA, because the evolved Δ*ampC* decreased in growth rate and both Δ*ampC* and TCR reduced resistance during the reverse evolution (Fig.3c), even though the Δ*ampC* and TCR acquired less resistance than WT and CompA.

**Fig 3.**
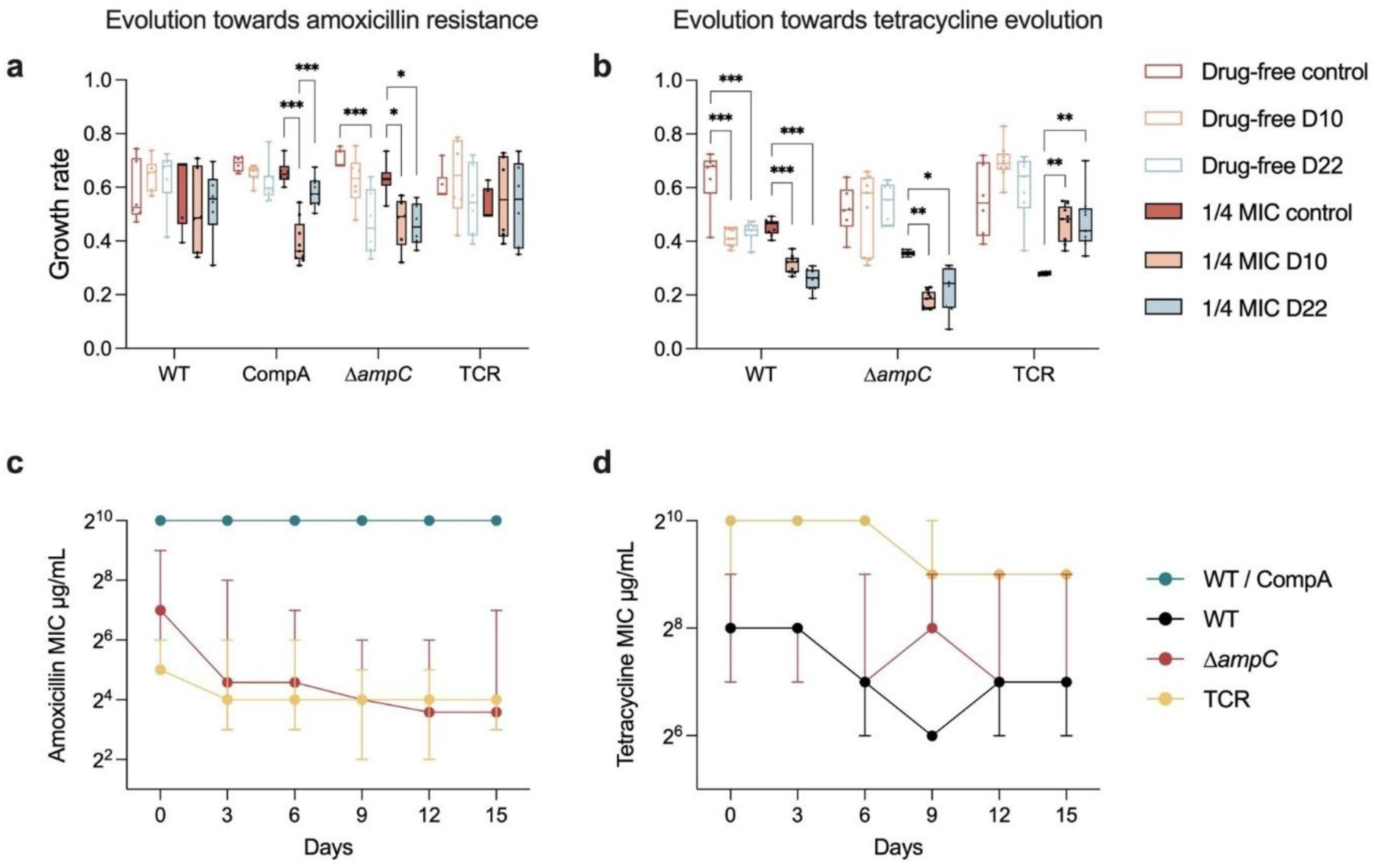
The fitness of resistant strains. Growth rate of amoxicillin (**a**) or tetracycline (**b**) resistant strains at day 10, day 22 and naive control in the evolution experiments in drug-free or ¼ MIC condition. The significance was calculated by ANOVA and shown with compact letter display. The significance of the differences between groups of the same strain is indicated with asterixis: \*\*\**p*<0.001, ** *p*<0.01, * *p*<0.05. **c** and **d** show the resistance change during the reverse evolution in drug-free medium after the amoxicillin (c) or tetracycline (d) resistance evolution.

During the tetracycline resistance evolution, the growth rate of TCR and Δ*ampC* did not change in the drug-free condition, while that of WT decreased in this condition (Fig.3b). At ¼ MIC condition, unlike the growth rate of WT and Δ*ampC,* which decreased at day 10 and day 22, that of TCR increased significantly during resistance evolution. The tetracycline resistance of all evolved strains was reduced in 15-day reverse evolution (Fig.3d). The decrease of resistance in WT and Δ*ampC* was similar to that of TCR. The combined data suggest a similar fitness burden of the mechanism of tetracycline resistance in all strains.

### Genome structure variation of evolved strains

To determine whether the GDA is still prevalent in *ampC* knocked out and replaced mutants, whole genome sequencing (WGS) was used for the identification of genome structure variation in amoxicillin and tetracycline evolved strains. The *ampC* gene in WT and the CompA and *tet(B)* genes in TCR are resistance genes for amoxicillin and tetracycline, respectively. The *ampC* GDA were found in all amoxicillin evolved WT and CompA strains. The *tet(B)* GDA was only observed in 5 of 9 tetracycline evolved TCR (Fig.4).

**Fig. 4.**
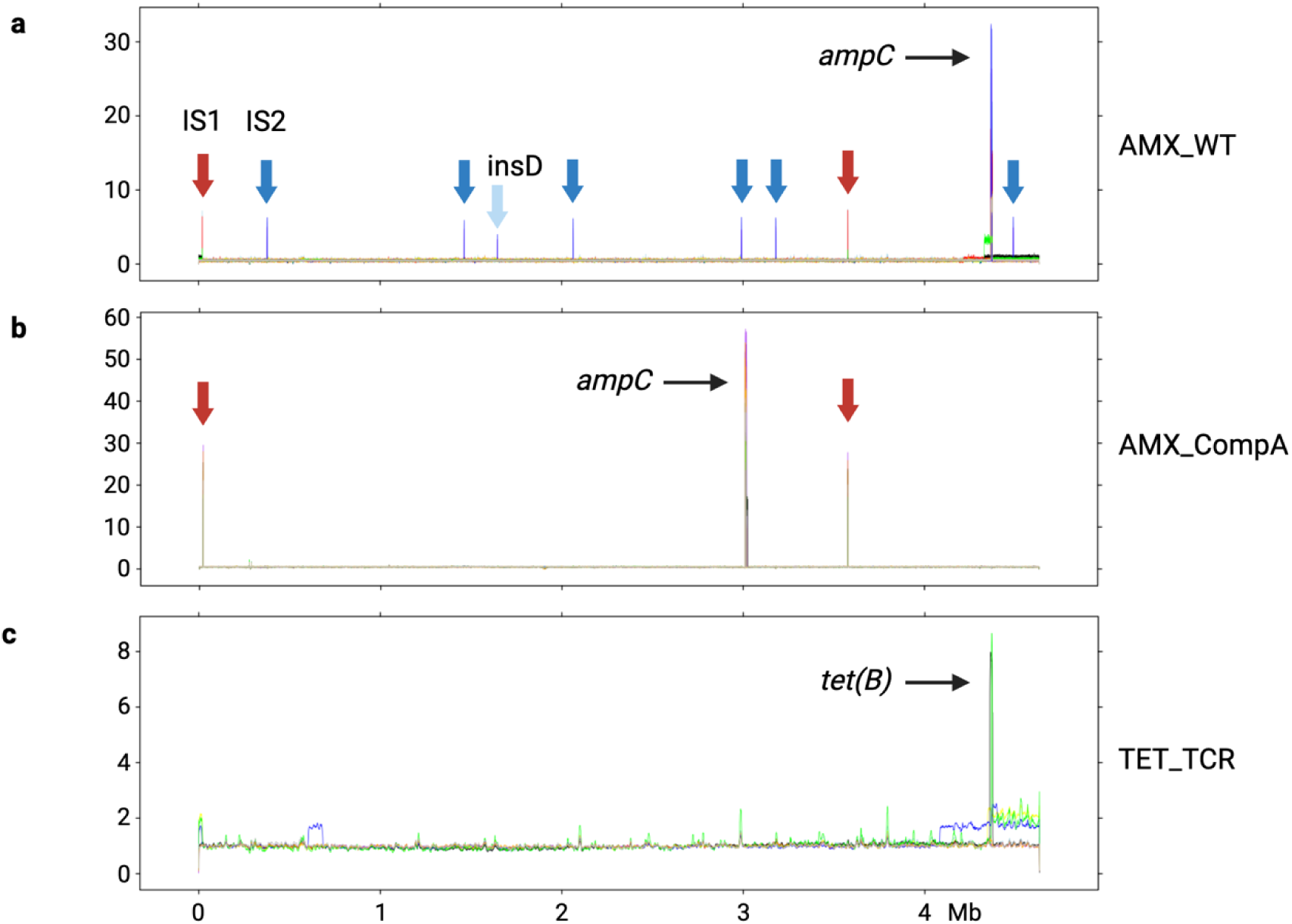
The GDA in the evolved strains with an efficient resistance gene. Panels: (**a**) *de novo* amoxicillin resistant WT, (**b**) *de novo* amoxicillin resistant CompA and (**c**) *de novo* tetracycline resistant TCR. The X axis is the genome-wide position in *E. coli*. The Y axis is the normalized read counts of WGS. Note that the scales on the Y-axes differ.

Genomic DNA is a supercoiled loop. Several large duplicons were observed at the start and the end of reference genome (Fig.4ac). The identified duplicons tend to exhibit larger size than amplicons. Three duplicons were observed, one in amoxicillin evolved WT and two in tetracycline evolved TCR (Fig.4ac). The size varied from 32.69 to 323.57 kb (Fig.5a), located between 4.33 to 0.02 Mb in the genome. With the exception of one amplicon in an amoxicillin evolved WT with 7.27 copies that had a large size, 46.42 kb, located between 33 and 4.37 Mb in the genome, the size of all other amplicons ranged from 2.94 to 16.41 kb (Fig.5a).

**Fig. 5.**
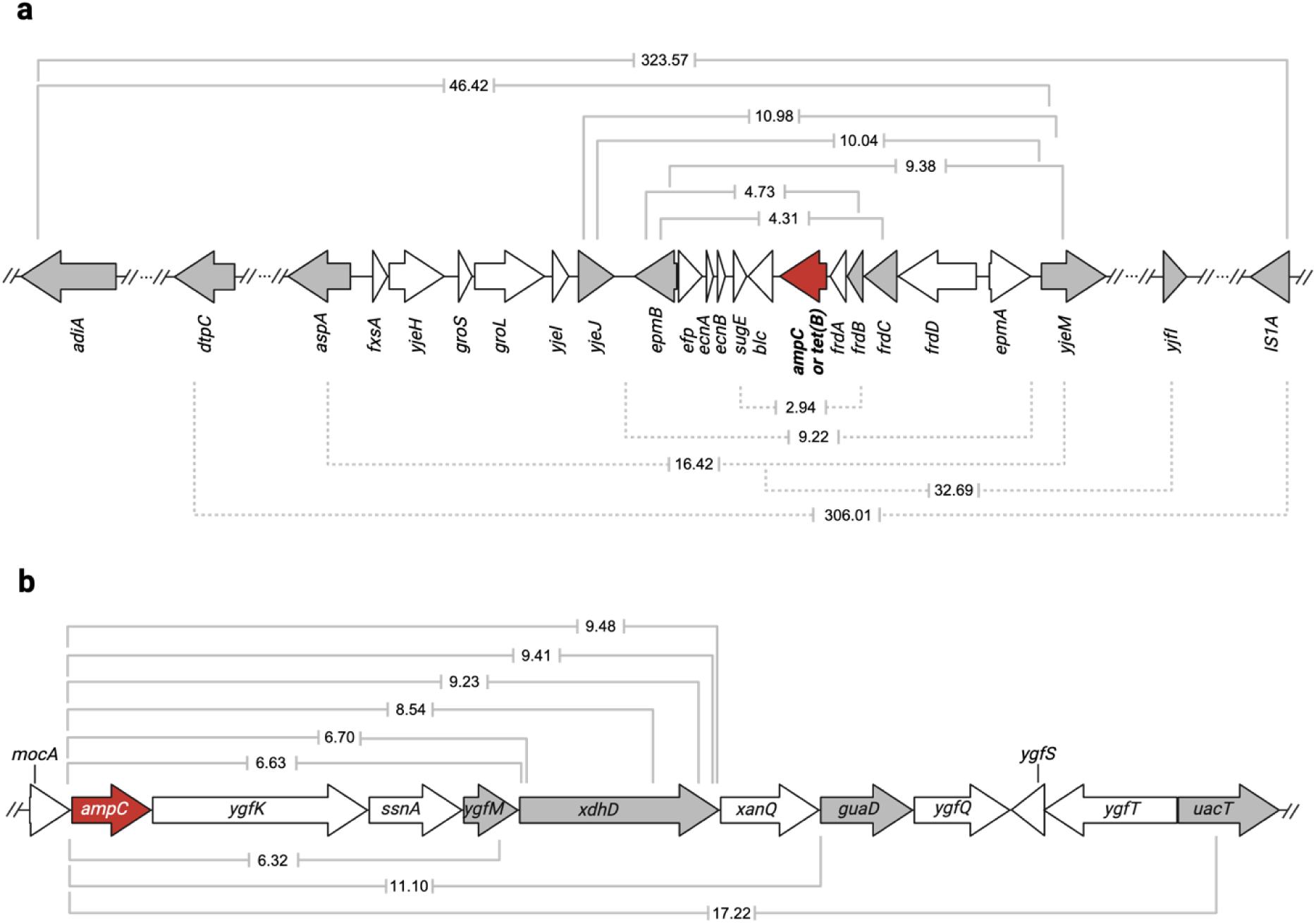
Amplicons or duplicons in the evolved strains containing a resistance gene. The size (kb) is indicated in the brackets. **a** The amplicons or duplicons of *de novo* amoxicillin resistant WT (solid brackets) and *de novo* tetracycline resistant TCR (dotted brackets). **b** The amplicon of *de novo* amoxicillin resistant CompA is shown using solid brackets. The resistance gene is indicated in red. The gene cassettes in which fragments start or end are shown in grey.

The junction of GDAs reveals some information about the mechanism of amplification (19). There are two reported kinds of junction for *ampC* amplification in *E. coli*, one involves transposon element IS1 (*insA* and *insB* genes) and the second entails direct tandem connection (17). The sequence of the GDA junction can be confirmed by the computational pipeline Breseq based on WGS raw data. Moreover, if the amplification mechanism involves a transposon element, the read counts of amplicon and transposon element in WGS is higher than the median of the reference genome.

In the sequenced amoxicillin evolved WT in this study, three out of nine replicates exhibit IS1 inserted into the GDA junction. The IS1 copies of these also show a genome wide increase (Fig.4a). The other six amoxicillin evolved WT have microhomology repeat sequences in their GDA junction without IS1 insertion. Four of these remained without any transposon elements frequency increase. Transposon element IS2 (*insC* and *insD* genes) was inserted into the amplicon, in the P*_ampC_* (Fig.6a), in two replicates. The IS2 copy increase was observed all over the genome. In evolved CompA, an increase in the read counts of IS1 and the fragment around the *ampC* gene was observed (Fig.4b), suggesting the *ampC* amplification in CompA could also be mediated by IS1. The copy number of *ampC* gene, calculated through normalized read counts, in evolved CompA was higher than that of evolved WT (Fig.4ab), although they exhibited similar amoxicillin MIC (Fig.1b).

**Fig. 6.**
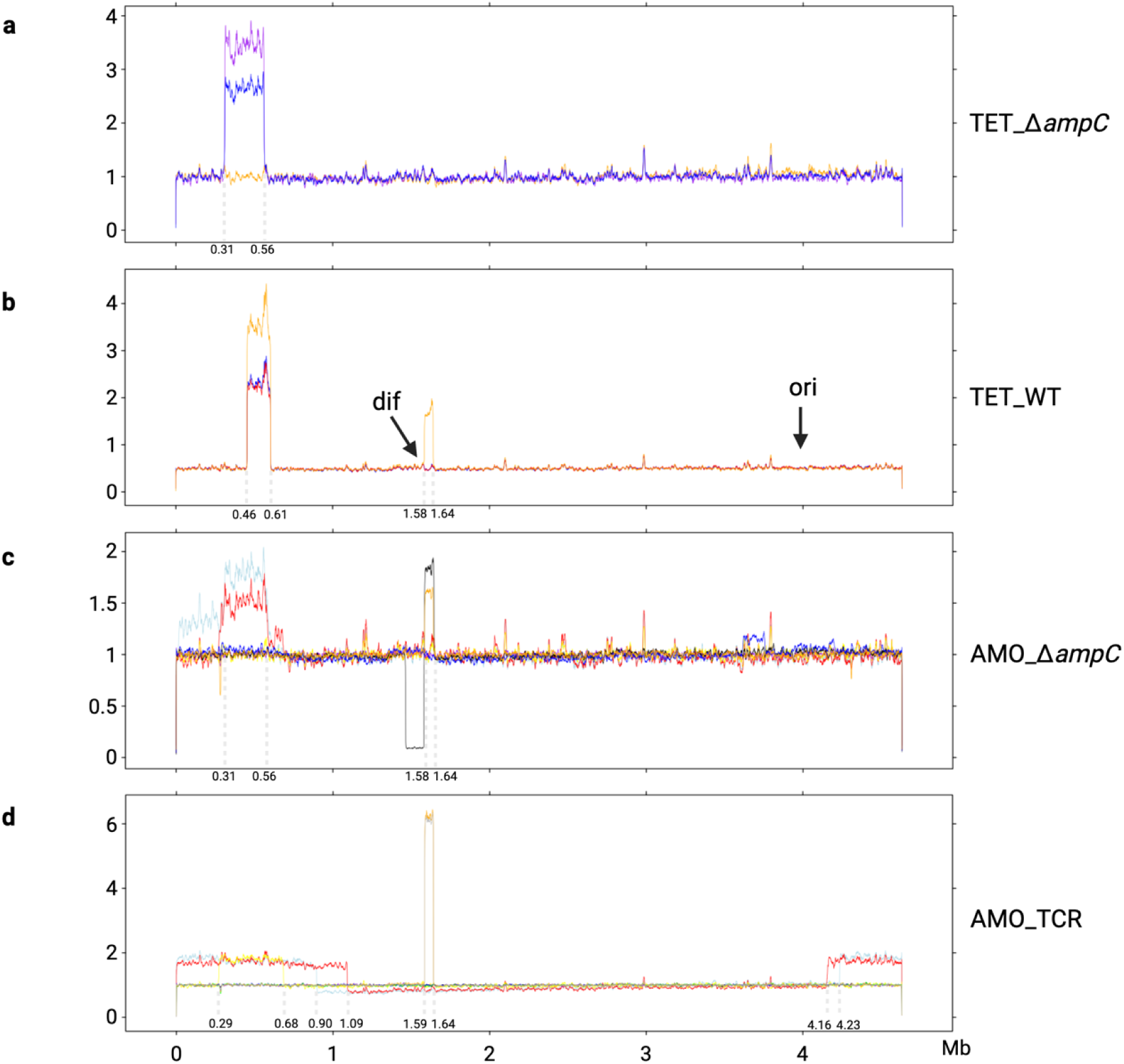
The read frequency variation after resistance evolution in WGS of strains lacking an effective resistance gene. (**a**) tetracycline resistant Δ*ampC* and (**b**) WT. The read frequencies of amoxicillin resistant Δ*ampC* and TCR are presented in panels **c** and **d**. Note that the scales of the Y-axes are not identical.

After the tetracycline resistance evolution, not all evolved TCR replicates contained the *tet(B)* gene amplification (Fig.4c), although their tetracycline MIC’s were similar. Five replicates exhibited the amplification and four did not. The highest copy number of *tet(B)* was around 8. Compared to the copy number after amoxicillin resistance evolution of the amplicon containing *ampC* in WT (around 30) and CompA (approximately 50) this is considerably lower. In one tetracycline evolved TCR replicate there was, in addition to the GDA of *tet(B)*, a duplication located in 0.6 to 0.7 Mb (Fig 4c). Several replicates had a wide duplicated shoulder next to the amplified *tet(B)*.

The amplicons and duplicons were different but comparable in amoxicillin evolved WT and tetracycline evolved TCR (Fig.5a). The largest size of a duplicon was 323.57 kb, shortest one was only 2.94 kb. Four out of 12 identified GDAs start at the *epmB* cassette, while six end at *yjeM* cassette. One duplicon of amoxicillin evolved WT and one of tetracycline evolved TCR ends at IS1A, which is located from bp 19,796 to bp 20,563 in genome.

The size of *ampC* amplicon in *de novo* amoxicillin resistant CompA ranged from 6.32 to 17.22 kb (Fig.5b). All CompA amplicons started at the same position in the *ampC* promoter and ended at different sites. Six out of nine amplicons ended at variable locations within the *xdhD* gene. This indicates that while a preferred starting point exists for the amplification, the end of the fragment can vary.

In some of the strains lacking an efficient resistance gene, amplification still occurred, but in a more diverse manner and involving larger fragments of the chromosomal DNA (Fig.6). Three kinds of GDA events were discovered, located at 0.29 to 0.68 Mb, 1.58 to 1.64 Mb, 4.16 to 1.09 Mb. The selection of the GDA without an efficient resistance gene occurred less often and may carry more fitness costs. Not all replicates of these resistant strains generated a GDA event. Only one tetracycline resistant WT replicate contained the GDA of 0.29 to 0.68 Mb and 1.58 to 1.64 Mb at the same time. The GDA of 0.29 to 0.68 Mb and 1.58 to 1.64 Mb occurred in both amoxicillin and tetracycline resistant strains, indicating that the mechanism for generating these GDAs consists of a set of genes causing multidrug resistance. Two amoxicillin resistant TCR replicates exhibited a large fragment duplication (4.16 to 1.09 Mb). This large fragment contained both the other identified GDA fragment from 0.29 to 0.68 Mb and the *tet(B)* gene located at 4.37 Mb. The amoxicillin resistant TCR generated a larger GDA fragment than the amoxicillin resistant Δ*ampC*.

The genomic DNA replication starts at the *ori* site (232 bp), finishes and separates at the *dif* site (28 bp) (27).The large duplicon (4.16 to 1.09 Mb) in amoxicillin resistant TCR does not cross the *ori* and *dif* sites. The start of the GDA from 1.6 to 1.7 Mb is located at the *dif* site and the ends at a copy of *insD*. In one amoxicillin resistant Δ*ampC*, the duplication of 1.58 to 1.64 Mb led to a deletion between 1.4 to 1.6 Mb. This suggests this duplication might be caused by the exchange of genomic replicons.

### Promoter mutations of *de novo* resistant genes

Mutations in promoter areas are a main cause of *ampC* transcription increase in beta-lactam resistant strains (28). The accumulation of mutations in the promoter of *ampC* (P*_ampC_*) occurred before *ampC* amplification during amoxicillin resistance evolution in WT *E. coli* (17). Unlike GDA, which causes overexpression of other hitchhiking genes near the resistance gene, the mutations in gene promoters exert specific impact. In order to investigate the frequency of the mutations in P*_ampC_* when it regulates none or different resistance genes, the P*_ampC_* of all resistant strains was cloned by PCR and subjected to Sanger sequencing (Fig 7).

**Fig. 7.**
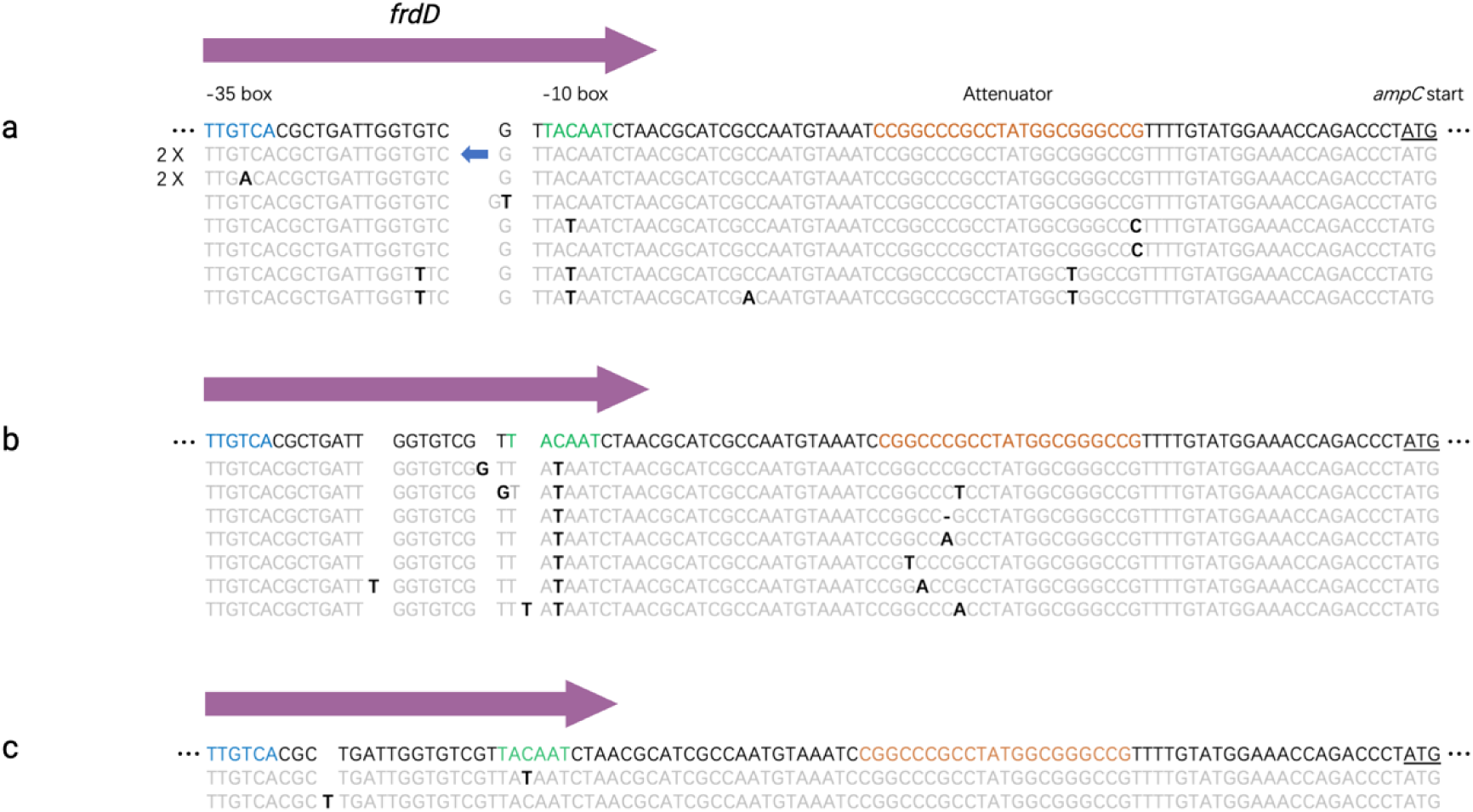
The promoter mutations in resistant strains. **(a)** The P*_ampC_* mutations of amoxicillin resistant WT. **(b)** The P*_ampC_* mutations of tetracycline resistant TCR. **(c)** The P*_ampC_* mutations of amoxicillin resistant CompA. The first line of each figure represents the sequence of the original P*_ampC_*without any mutations. The promoter elements including the −35 box, the −10 box and the attenuator are indicated in blue, green and orange, respectively. The reading frame of *frdD* is marked with a purple arrow. The *ampC* transcriptional start site is underlined. The point mutations are indicated by black and bold letters. Unmutated bases are in grey. The blue arrow in the upper sequence indicates IS2.

There are 3 elements that impact the strength of P*_ampC_*. These are the RNA polymerase RpoD binding sites in the −10 box and the −35 box, and thirdly an attenuator (29). The −10 box and - 35 box are within the reading frame of *frdD* gene, which is next to *ampC* gene. Thus, the mutations in P*_ampC_* may impact the function of FrdD. Promoter mutations were observed in all amoxicillin evolved WT (Fig.7a). 6 out of 9 replicates of amoxicillin evolved WT contained mutations in these three elements. An IS2 transposon was identified inserted between the −10 box and −35 box in two amoxicillin evolved WT populations. This insertion may inactivate the *frdD* gene and impact the strength of P*_ampC_*. The whole *ampC* cassette is in the amplicon. The consequence is that the number of IS2 copies increased in these two replicates throughout the genome (Fig.4a).

Although only half of tetracycline evolved TCR replicates exhibited the GDA containing the *tet(B)* gene, most replicates acquired the promoter mutations. 7 out of 9 tetracycline evolved TCR acquired the P*_ampC_* mutations, which are all related to the −10 box and the attenuator (Fig.7b). Only 2 out of 9 amoxicillin evolved CompA acquired the P*_ampC_*mutations, even though the promoter was translocated together with the *ampC* gene (Fig.7c). One mutation was identified in −10 box and the other one was an insertion base between −10 box and −35 box. The IS1 insertion with both directions was detected in 8 of 9 resistant CompA at 34 to 38 bp before the −35 box. No mutations occurred in the P*_ampC_* of amoxicillin evolved Δ*ampC* and TCR (data not shown). The absence of point mutations in Δ*ampC* indicates that the presence of *ampC* is essential for the P*_ampC_* mutations to occur. As the *tet(B)* slightly contributes amoxicillin resistance, the mutations in the promoter might also enhance resistance. However, promoter mutations were not detected in amoxicillin evolved TCR.

### Point mutations in the evolved strains

To identify the point mutations of the evolved strains with different initial genomic backgrounds, the genome of all replicate strains was sequenced after resistance evolution. The DNA was isolated from a population after evolution rather than from a single colony. The allele frequency of point mutations in the sequencing was used to describe the landscape of resistance variants’ genotype. As the genomic mutations with high frequency have more potential to impact the resistance, the point mutations with frequency lower than 0.1 or also appearing in the control grown in the absence of the antibiotic were removed from the analysis.

In total, there are 75, 25, 120 and 81 point mutations detected in the amoxicillin resistant WT, CompA, Δ*ampC* and TCR, respectively (Supplemental Table 1-7). The point mutations related to cell membrane, transcription, DNA repair or stress response and other essential genes are presented in the heatmap in Fig.8. Although all evolved WT and CompA replicates acquired the same amoxicillin resistance after evolution, more point mutations accumulated in evolved WT than in CompA. Without the *ampC* gene, in evolved Δ*ampC* and TCR, more point mutations accumulated. Compared to WT and CompA, the mutations in amoxicillin target *ftsI* and efflux pump *acrB* were more common in Δ*ampC* and TCR. The evolved Δ*ampC* exhibited more point mutations than TCR.

**Fig. 8.**
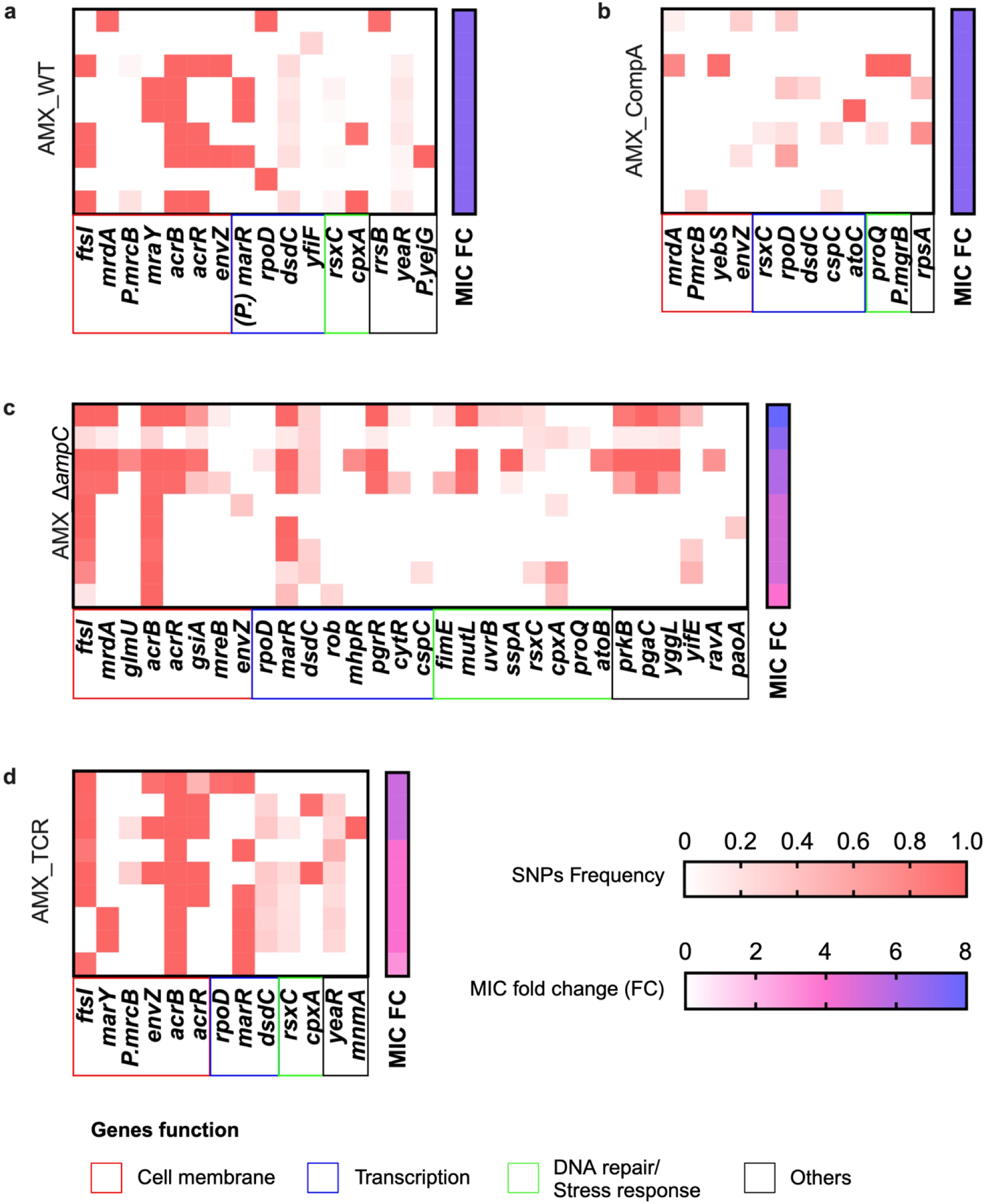
Frequency of mutated gene in amoxicillin 22-day evolved replicates of **(a)** WT, **(b)** CompA, **(c)** Δ*ampC* and **(d)** TCR. Each row represents a sequenced independent replicate of the evolution experiments. Fold change of MIC by the power 2 of the sequenced population is shown on the right of each heatmap. Genes are grouped by function.

After 22-day tetracycline resistance evolution, 204, 31 and 46 point mutations were identified in the evolved TCR, WT and Δ*ampC* replicates, respectively. The number of mutations of resistant TCR was considerably higher than that of resistant WT and Δ*ampC*. The most common mutated genes in TCR were efflux pump *acrR*, and transcriptional regulators *marR* and *rob*. Many mutated genes are involved in metabolic pathways, such as fatty acid metabolism (*fabA*, *fabB*, *fabD*, *fadE*, *fabG*, *fadR*) and secondary metabolites biosynthesis (*icd*, *speC*, *purF*, *glyA*, *pck*, *gcvH*, *galM*) (Fig.9a). In the WT and Δ*ampC*, the common mutated genes were membrane sensor histidine kinase *envZ*, and trancriptional regulator *ybaO* (Fig.9b). The *rob* mutation was not observed in resistant WT and Δ*ampC*, while the mutation in *ybaO* was not observed in resistant TCR.

**Fig. 9.**
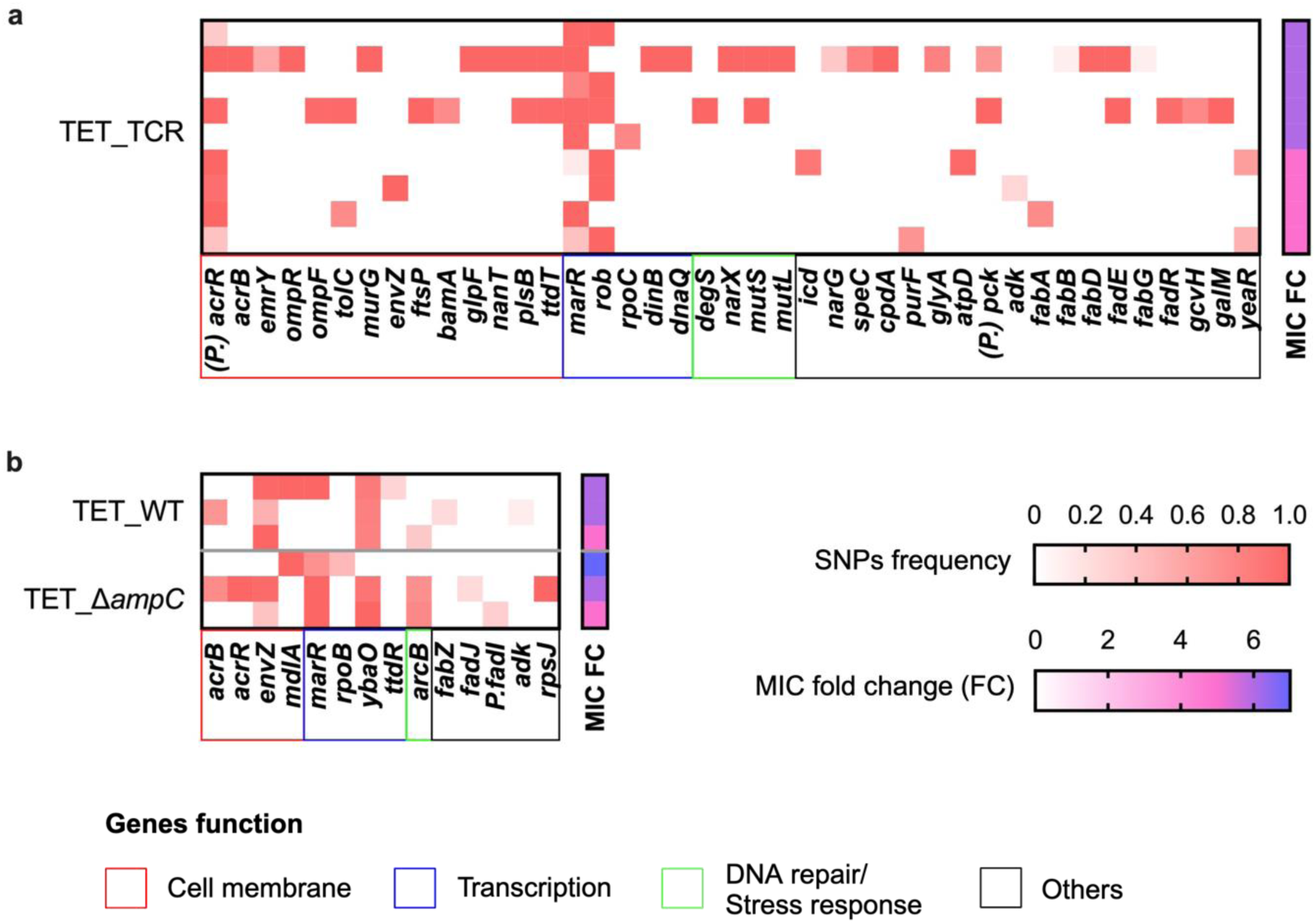
Frequency of mutated genes in tetracycline evolved **(a)** TCR, **(b)** WT and Δ*ampC*. Each row represents the sequenced populations from replicates of independent evolution experiments. Fold change of MIC by the power 2 of the sequenced population are shown on the right of each heatmap. Genes are grouped by function.

## Discussion

Amplification of fragments of chromosomal DNA containing resistance genes is one of the mechanisms by which bacteria can develop high levels of resistance against antibiotics (19, 30, 31). In the case of induced amoxicillin resistance in *E. coli*, amplification with fragment containing *ampC* gene is a common event (4). When the *ampC* gene was deleted, the fragment in that position was not be amplified under amoxicillin resistance evolution (17). Apparently, the cell is able to amplify a fragment containing a gene that codes for a protein needed at high expression levels to combat toxic growth conditions. If that is indeed correct, the following questions present themselves: What if the *ampC* gene is inserted at another location in the chromosome? Is then a comparable fragment around it amplified? When a weak resistance gene for another, very different, antibiotic is inserted into the location of the *ampC* gene, does the induction of resistance against that gene cause amplification of the same fragment?

When a strain with the *ampC* gene at another location (CompA) is induced to develop amoxicillin resistance, a similarly sized fragment is amplified around the new location, while the fragment around the original *ampC* gene location is not amplified. This indicates that the cell sets in motion the amplification of the fragment around *ampC* as a result of increased *ampC* expression. The *ampC* amplicons from CompA are more conserved than those of the WT, as all amplicons in CompA start at the same position and most of them end within the same gene. The start site of CompA amplicon is upstream of the *ampC* promoter. This site was translocated together with the *ampC* gene when CompA was constructed. Although this site has the same distance to the *ampC* gene in the WT, the amplification was not observed to start at this site in amoxicillin resistant WT in this or other comparable studies (4, 9, 17, 18, 32). This indicates that the position of the resistance gene does impact the size and boundaries of the amplified fragments.

GDA occurred in some evolved TCR replicates, in which the *ampC* gene is replaced by the *tet(B)* gene. The GDAs of TCR end within several genes (*frdC*, *yjeM*, *IS1A*) which were also observed in amoxicillin resistant WT. This suggests that the cell utilized a similar mechanism to cope with the tetracycline challenge. The duplication fragment with *tet(B)* in amoxicillin resistant TCR was much larger (from 4.16 to 1.09 Mb), indicating that at least two resistance causing mechanisms exist in parallel, giving comparable, but not identical results. In both cases, the cell amplified a DNA fragment in response to the demand for higher expression levels of a resistance gene. As a result, cells that increase expression levels of a resistance gene through amplification of a chromosomal DNA fragment have higher fitness under exposure to the corresponding antibiotic.

AmpC inactivates the antibiotic and Tet(B) actively pumps out the drug, both causing resistance. These mechanisms carry different fitness costs. The AmpC protein is a periplasmic enzyme, not known to cause any cellular toxicity and does not consume ATP when breaking the beta-lactam ring (33). The efflux pump Tet(B), however, is dependent on energy through proton exchange to reduce the intracellular drug concentration (34). Overexpression of the *ampC* and *tet(B)* genes conveys a selection advantage at different levels. All amoxicillin evolved WT and CompA relied on the overexpression of *ampC*, through GDA and promoter mutations, for resistance development. However, not all tetracycline resistant replicates showed amplification of the *tet(B)* gene. The long-term compensatory evolution can lead to reduction of resistance through fitness selection or additional fitness-compensatory mutations maintaining resistance (35). The loss of tetracycline resistance in evolved TCR during the compensatory evolution, suggests that the energy consumption by the mechanism for resistance impacts resistance maintenance.

The duplications observed in the *ampC* knock-out located between 0.29 to 0.68 Mb or 1.58 to 1.64 Mb, indicate the presence of another resistance gene there. The resistance gene between 0.29 to 0.68 Mb could be *acrAB,* coding for a multidrug efflux pump. AcrAB is responsible for the resistance of many drugs including tetracycline, ampicillin, chloramphenicol and several others (36). The resistance gene located between 1.58 to 1.64 Mb could be the multiple antibiotic resistance family (Mar) (37). The *marRAB* operon participates in controlling the transcription of resistance genes and thus facilitates the development of antibiotic resistance (38). This operon is implicated in the regulation of at least 80 genes involved in multidrug efflux (39), oxidative stress (40), biofilm formation (41) and other processes (42). The sizes of the duplicon and amplicon at these locations are larger than that of *ampC* and *tet(B)* in amoxicillin and tetracycline resistance strains. This explains the lower copy number of the fragments at 0.29 to 0.68 Mb and at 1.58 to 1.64 Mb, as the higher copy number of the *ampC* fragment is associated with a narrower amplified region (3). Low fitness cost of amplification is necessary for its enduring occurrence (30). The multidrug resistance systems are multi-targeting. The activation of these systems has greater fitness consequences, which may be ameliorated by the overexpression of other regulons. In conclusion, the cellular systems involved in development of high levels of *de novo* resistance achieve this resistance at minimal fitness costs due to their high specificity.

The accumulation of GDA and point mutations influence each other. Similar to *ampC* GDA, the mutations in the promoter can also significantly increase the transcription of *ampC* (17, 29, 43, 44). Fewer mutations were accumulated in amoxicillin resistant CompA. Thus, more *ampC* gene copies are needed to compensate for the lower mutation frequency at P*_ampC_*.

The observation that amoxicillin resistant strains with *ampC* gene, WT and CompA, had fewer point mutations than Δ*ampC* and TCR suggests that the *ampC* GDA can alleviate the necessity of some mutations in resistance evolution. Similar amplifications of the efflux pump gene *sdrM* bypasses multiple target mutations against delafloxacin resistance in *Staphylococcus aureus* (15). The *ampC* amplification bypasses the single-nucleotide polymorphism (SNPs) in *acrB*/*ftsI*/*envZ* in *E. coli* (17). Additionally, the GDA of *acrAB* or *mar* family is not accumulated in amoxicillin resistant WT and CompA. This suggests that the *ampC* amplification prevented GDA in other locations during beta-lactam adaptation by making these unnecessary.

Compared to amoxicillin resistance, tetracycline resistant TCR had significantly more point mutations, especially in genes related to metabolic pathways. Efflux pumps are located in the cell membrane (45). The overproduction of Tet(B) may disrupt the synthesis of membrane components, leading to more mutations in fatty acid metabolic pathways in *de novo* tetracycline resistant strains.

The mechanisms of GDA can be classified as RecA-dependent or RecA-independent (19). However, it should be noted that several RecA-independent mechanisms to some extent still rely on RecA-mediated homologous recombination (10). The repetitive sequence at the junction of GDA is used to distinguish these two manners. If the GDA arises between longer repeat sequences, in the order of 20-40 nucleotides, the GDA is mediated by a RecA-dependent mechanism, otherwise, by a RecA-independent system (19, 46). When GDA has insertion sequences in their junctions, the GDA mostly is based on a RecA-dependent mechanism, as the identity of two insertion sequences is used as homology region for recombination (47). However, there is evidence that the GDA related to insertion sequences is transposition-dependent and RecA-independent in some situations (23, 24). The *ampC* amplification is usually determined by means of RecA-independent mechanisms as the GDA can form a short direct repeat in the junction (12-13 bp) and still occur in *recA* inactive mutants (19, 48). However, several amoxicillin evolved strains contained transposon element IS1 in their *ampC* amplification junctions, which indicates that the GDA can also be mediated through a RecA-dependent or transposition-dependent mechanism (17). Two duplicons were identified to end at IS1A, which is 284 kb away from the resistance gene *ampC* or *tet(B)* in the genome. When the copy number increases, the amplicons’ size narrow, while the IS1 remained in the amplification junction. This indicates that the functional IS1 may be involved in the GDA events.

The GDA junctions between 0.29 to 0.68 Mb, 1.58 to 1.64 Mb, 4.16 to 1.09 Mb exhibit 3-5 bp short repeat sequences, indicating a RecA-independent duplication mechanism. This suggests their occurrence is largely independent of homology recombination (49). The GDA may occur during the DNA replication process, as the GDA between 1.58 and 1.64 Mb starts at a 28 bp *dif* site which is the separation location of newly replicated bacterial chromosomes (27). The amplification of *dif* site may in turn impact the DNA replication process. Even when the GDA fragment is very large, increasing to a quarter of the chromosome size (4.16 to 1.09 Mb), it does not cross the DNA replication origin and segregation position. This suggests the mechanism of the GDA probably operates through exchange with the sister-strand (50).

Regarding the hypothesis formulated in the introduction, the main conclusions are:

1. Amplification occurs independent of the location of the gene within the chromosome.
2. Amplification of effective resistance genes prevents GDA at another position.
3. Amplification is not limited to a single specific gene.
4. The fitness costs resulting from the resistance gene affect the copy number of the GDAs.
5. The amplicon or duplicon does not cross the DNA replication origin and separation site.

On a speculative note, one could state that the induced amplification of a chromosomal fragment containing a gene that codes for a product that the cell needs in higher quantities than usual, is an inversion of the normal flow of information. Instead of the usual DNA to RNA to protein flow, the increased need for the protein induces DNA editing and processing. It remains to be seen whether this is a one-of-kind rarity, or a process speeding up evolution that was up till now not recognized as such.

## Materials and Methods

### Stains, growth condition and antibiotic

The WT *E. coli* used in this study is BW25113. Δ*ampC* (JW4111, BW25113_Δ*ampC*) was purchased from Keio library (National BioResource Project) and the kanamycin resistance cassette was removed. The TCR and CompA mutants were constructed by λ Red-mediated recombination (51). Primers used in the study are shown in Supplemental Table 8. In the CompA strain construction, the *ampC* cassette, between 4,367,403 and 4,368,895 in *E. coli* BW25113 genome, was deleted. Then this cassette was inserted into 3,009,241 positions, between *mocA* and *ygfK* genes. The selection resistance cassette was removed using the FRT site and FLP recombinase. The tetracycline resistance gene *tet(B)* in TCR mutant was clone from *E. coli* BL21-Gold.

All strains were grown in LB liquid medium at 37 °C shaken at 200 rpm. Stock solutions of amoxicillin and tetracycline were made in concentrations of 10 mg/mL. The antibiotic stock was filter-sterilized and kept at 4 °C. New stocks were made every three days.

### Evolution

Before the resistance adaptation evolution, the strains were grown in LB liquid medium, and the MIC’s of the cultures were measured. Then the strains were passaged to fresh medium controlling the initial OD_600_ as 0.1 with ¼ MIC antibiotic, which is day 0 of the evolution. After 24 h culture, the strains were passaged to fresh medium with two different concentrations. One is the same as the previous day, the other one is two-fold higher. After 24 h incubation, the OD_600_ was measured. If the OD_600_ of the culture with the higher antibiotic concentration is more than 70 % of the OD_600_ of the culture with the lower concentration, the culture with the higher concentration was passaged to the same growing condition and another one with double the antibiotic concentration. Otherwise, the lower one was used. This passage was repeated for 22 days.

The reverse evolution lasted 15 days after the resistance evolution. The resistant strains were passaged in drug free LB medium with the same growing conditions described above.

### MIC measurement

The MIC values were measured by growing strains in medium containing a gradient dilution of antibiotic in 96-well plates. The dilution constant was 2. The initial OD_600_ of each well was 0.05. The 96-well plates were grown in plate readers (Thermo Scientific Multiskan FC with SkanIt software) with pulse shaking at 37 °C for 24 h. The measurements of OD_595_ were conducted every 10 min. The MIC was defined as the lowest antibiotic concentration that restrains growth to OD_600_ less than 0.2 after 24 h incubation.

### Growth rate measurement

The growth rate was calculated from the log phase in the growth curve. The strains were cultured with 150 µL drug-free or ¼ MIC antibiotic liquid LB medium in round bottom 96- well plates in a plate reader at 37°C with pulse shaking for 24 h. The initial OD_600_ was 0.05. The OD_600_ was measured every 10 min. For each strain at least three replicates were used. The growth rate was calculated in R with the Growthrates package (52).

### Promoter sequencing

The *ampC* promoter was cloned by polymerase chain reaction (PCR) with PrimeSTAR Max (TaKaRa) as DNA Polymerase, isolated genomic DNA as template, F-AmpC Prom and R- AmpC prom (Supplemental Table S8) as primers. The PCR product was purified with MSBSpinRapace kit (Stratec). The purified product was then sequenced by Sangar (Macrogen Europe) and analyzed through Snapgene.

### Whole-genome sequencing

Illumina (NextSeq 550 system) was utilized for sequencing the DNA isolated from 1 mL culture. The DNA library preparation was performed using established protocols (13). After sequencing the reads were aligned to the reference CP009273, and analyzed for genetic and structural variation as described in Nong et al (13). The single nucleotide polymorphisms with allele frequency lower than 0.1 and those also present in the control were removed.

### Statistical analysis

The resistance adaptation speed between different strains was compared using as statistical analysis the Wilcoxon signed-rank text in R (53). The average of all replicates at each time point in each strain was calculated. The significance was determined by the p value calculated between these data sets.

The one-way ANOVA was used for the statistical significance determination of the MIC and growth rate between strains. The log_2_ transformed MIC was used in analysis to eliminate the influence of the dilution constant.

## Author contributions

LN performed research, designed experiments and wrote the first draft; XL assisted with data analysis; XW assisted with experimentation, WdL performed bioinformatic analysis, MJ assisted with experimental design and genomic analysis; SB assisted with data interpretation; BtK assisted with experimental designed, supervised the project and co-wrote the manuscript. All authors have commented on an earlier version of the manuscript and approve of the final draft.

## Data availability

The data that support the findings of this study are available from the corresponding author upon request.

## References

1. Poudel AN, Zhu S, Cooper N, Little P, Tarrant C, Hickman M, Yao G. 2023. The economic burden of antibiotic resistance: A systematic review and meta-analysis. PLOS ONE 18:e0285170.

2. Blaser MJ. 2016. Antibiotic use and its consequences for the normal microbiome. Science 352:544–545.

3. Gross R, Yelin I, Lázár V, Datta MS, Kishony R. 2024. Beta-lactamase dependent and independent evolutionary paths to high-level ampicillin resistance. Nat Commun 15:5383.

4. Darphorn TS, Hu Y, Sintanneland BBK, Brul S, Kuile BH ter. 2021. Multiplication of ampC upon Exposure to a Beta-Lactam Antibiotic Results in a Transferable Transposon in Escherichia coli. Int J Mol Sci 22:9230.

5. Gerland U, Hwa T. 2009. Evolutionary selection between alternative modes of gene regulation. Proc Natl Acad Sci 106:8841–8846.

6. Hoeksema M, Jonker MJ, Brul S, Kuile BH ter. 2019. Effects of a previously selected antibiotic resistance on mutations acquired during development of a second resistance in Escherichia coli. Bmc Genomics 20:284.

7. Händel N, Schuurmans JM, Brul S, Kuile BH ter. 2013. Compensation of the Metabolic Costs of Antibiotic Resistance by Physiological Adaptation in Escherichia coli. Antimicrob Agents Ch 57:3752–3762.

8. Händel N, Schuurmans JM, Feng Y, Brul S, Kuile BH ter. 2014. Interaction between Mutations and Regulation of Gene Expression during Development of De Novo Antibiotic Resistance. Antimicrob Agents Chemother 58:4371–4379.

9. Hoeksema M, Jonker MJ, Bel K, Brul S, Kuile BH ter. 2018. Genome rearrangements in Escherichia coli during de novo acquisition of resistance to a single antibiotic or two antibiotics successively. Bmc Genomics 19:973.

10. Reams AB, Kofoid E, Savageau M, Roth JR. 2010. Duplication Frequency in a Population of Salmonella enterica Rapidly Approaches Steady State With or Without Recombination. Genetics 184:1077–1094.

11. Pettersson ME, Sun S, Andersson DI, Berg OG. 2008. Evolution of new gene functions: simulation and analysis of the amplification model. Genetica 135:309.

12. Tomanek I, Grah R, Lagator M, Andersson AMC, Bollback JP, Tkačik G, Guet CC. 2020. Gene amplification as a form of population-level gene expression regulation. Nat Ecol Evol 4:612–625.

13. Nong L, Jonker M, Leeuw W de, Wortel MT, Kuile B ter. 2024. Progression of ampC amplification during de novo amoxicillin resistance development in E. coli. mBio e0298224.

14. Tomanek I, Guet CC. 2022. Adaptation dynamics between copy-number and point mutations. eLife 11:e82240.

15. Silva KPT, Sundar G, Khare A. 2023. Efflux pump gene amplifications bypass necessity of multiple target mutations for resistance against dual-targeting antibiotic. Nat Commun 14:3402.

16. Qi W, Jonker MJ, Leeuw W de, Brul S, Kuile BH ter. 2023. Reactive oxygen species accelerate de novo acquisition of antibiotic resistance in E. coli. iScience 26:108373.

17. Nong L, Jonker M, Leeuw W de, Wortel MT, Kuile B ter. 2024. Progression of ampC amplification during de novo amoxicillin resistance development in E. coli. bioRxiv 2024.05.24.595737.

18. Qi W, Jonker MJ, Leeuw W de, Brul S, Kuile BH ter. 2024. Role of RelA-synthesized (p)ppGpp and ROS-induced mutagenesis in de novo acquisition of antibiotic resistance in E. coli. iScience 27:109579.

19. Sandegren L, Andersson DI. 2009. Bacterial gene amplification: implications for the evolution of antibiotic resistance. Nat Rev Microbiol 7:578–588.

20. Tang S, Conte V, Zhang DJ, Žedaveinytė R, Lampe GD, Wiegand T, Tang LC, Wang M, Walker MWG, George JT, Berchowitz LE, Jovanovic M, Sternberg SH. 2024. De novo gene synthesis by an antiviral reverse transcriptase. Science 386:eadq0876.

21. Maybin M, Ranade AM, Schombel U, Gisch N, Mamat U, Meredith TC. 2024. IS1- mediated chromosomal amplification of the arn operon leads to polymyxin B resistance in Escherichia coli B strains. mBio 15:e00634–24.

22. Wei D-W, Wong N-K, Song Y, Zhang G, Wang C, Li J, Feng J. 2022. IS26 Veers Genomic Plasticity and Genetic Rearrangement toward Carbapenem Hyperresistance under Sublethal Antibiotics. mBio 13:e03340–21.

23. Harmer CJ, Hall RM. 2015. IS26-Mediated Precise Excision of the IS26-aphA1a Translocatable Unit. mBio 6:10.1128/mbio.01866-15.

24. Hubbard ATM, Mason J, Roberts P, Parry CM, Corless C, Aartsen J van, Howard A, Bulgasim I, Fraser AJ, Adams ER, Roberts AP, Edwards T. 2020. Piperacillin/tazobactam resistance in a clinical isolate of Escherichia coli due to IS26-mediated amplification of blaTEM-1B. Nat Commun 11:4915.

25. Darby EM, Trampari E, Siasat P, Gaya MS, Alav I, Webber MA, Blair JMA. 2022. Molecular mechanisms of antibiotic resistance revisited. Nat Rev Microbiol 1–16.

26. Akhtar AA, Turner DPJ. 2022. The role of bacterial ATP-binding cassette (ABC) transporters in pathogenesis and virulence: Therapeutic and vaccine potential. Microb Pathog 171:105734.

27. Castillo F, Benmohamed A, Szatmari G. 2017. Xer Site Specific Recombination: Double and Single Recombinase Systems. Front Microbiol 8:453.

28. Caroff N, Espaze E, Bérard I, Richet H, Reynaud A. 1999. Mutations in the ampC promoter of Escherichia coli isolates resistant to oxyiminocephalosporins without extended spectrum β-lactamase production. Fems Microbiol Lett 173:459–465.

29. Jaurin B, Grundström T, Normark S. 1982. Sequence elements determining ampC promoter strength in E. coli. Embo J 1:875–881.

30. Nicoloff H, Hjort K, Levin BR, Andersson DI. 2019. The high prevalence of antibiotic heteroresistance in pathogenic bacteria is mainly caused by gene amplification. Nat Microbiol 4:504–514.

31. Saathoff M, Kosol S, Semmler T, Tedin K, Dimos N, Kupke J, Seidel M, Ghazisaeedi F, Jonske MC, Wolf SA, Kuropka B, Czyszczoń W, Ghilarov D, Grätz S, Heddle JG, Loll B, Süssmuth RD, Fulde M. 2023. Gene amplifications cause high-level resistance against albicidin in gram-negative bacteria. PLOS Biol 21:e3002186.

32. Qi W, Jonker MJ, Katsavelis D, Leeuw W de, Wortel M, Kuile BH ter. 2024. The Effect of the Stringent Response and Oxidative Stress Response on Fitness Costs of De Novo Acquisition of Antibiotic Resistance. Int J Mol Sci 25:2582.

33. Knott-Hunziker V, Petursson S, Waley SG, Jaurin B, Grundström T. 1982. The acyl-enzyme mechanism of β -lactamase action. The evidence for class C β -lactamases. Biochem J 207:315–322.

34. Chopra I, Roberts M. 2001. Tetracycline Antibiotics: Mode of Action, Applications, Molecular Biology, and Epidemiology of Bacterial Resistance. Microbiol Mol Biol Rev 65:232–260.

35. Andersson DI, Hughes D. 2010. Antibiotic resistance and its cost: is it possible to reverse resistance? Nat Rev Microbiol 8:260–271.

36. Okusu H, Ma D, Nikaido H. 1996. AcrAB efflux pump plays a major role in the antibiotic resistance phenotype of Escherichia coli multiple-antibiotic-resistance (Mar) mutants. J Bacteriol 178:306–308.

37. Cohen SP, Hächler H, Levy SB. 1993. Genetic and functional analysis of the multiple antibiotic resistance (mar) locus in Escherichia coli. J Bacteriol 175:1484–1492.

38. Reyes-Fernández EZ, Schuldiner S. 2020. Acidification of Cytoplasm in Escherichia coli Provides a Strategy to Cope with Stress and Facilitates Development of Antibiotic Resistance. Sci Rep 10:9954.

39. Ruiz C, Levy SB. 2010. Many Chromosomal Genes Modulate MarA-Mediated Multidrug Resistance in Escherichia coli. Antimicrob Agents Chemother 54:2125–2134.

40. Alekshun MN, Levy SB. 1999. The mar regulon: multiple resistance to antibiotics and other toxic chemicals. Trends Microbiol 7:410–413.

41. Holden ER, Yasir M, Turner AK, Charles IG, Webber MA. 2022. Comparison of the genetic basis of biofilm formation between Salmonella Typhimurium and Escherichia coli. Microb Genom 8:mgen000885.

42. Barbosa TM, Levy SB. 2000. Differential Expression of over 60 Chromosomal Genes in Escherichia coli by Constitutive Expression of MarA. J Bacteriol 182:3467–3474.

43. Petrosino JF, Galhardo RS, Morales LD, Rosenberg SM. 2009. Stress-Induced β-Lactam Antibiotic Resistance Mutation and Sequences of Stationary-Phase Mutations in the Escherichia coli Chromosome▿. J Bacteriol 191:5881–5889.

44. Jaurin B, Grundström T. 1981. ampC cephalosporinase of Escherichia coli K-12 has a different evolutionary origin from that of beta-lactamases of the penicillinase type. Proc National Acad Sci 78:4897–4901.

45. Sun J, Deng Z, Yan A. 2014. Bacterial multidrug efflux pumps: Mechanisms, physiology and pharmacological exploitations. Biochem Biophys Res Commun 453:254–267.

46. Andersson DI, Hughes D. 2009. Gene Amplification and Adaptive Evolution in Bacteria. Annu Rev Genet 43:167–195.

47. Peterson BC, Rownd RH. 1983. Homologous sequences other than insertion elements can serve as recombination sites in plasmid drug resistance gene amplification. J Bacteriol 156:177–185.

48. Edlund T, Normark S. 1981. Recombination between short DNA homologies causes tandem duplication. Nature 292:269–271.

49. Reams AB, Roth JR. 2015. Mechanisms of Gene Duplication and Amplification. Cold Spring Harb Perspect Biol 7:a016592.

50. Lovett ST, Drapkin PT, Sutera VA, Gluckman-Peskind TJ. 1993. A sister-strand exchange mechanism for recA-independent deletion of repeated DNA sequences in Escherichia coli. Genetics 135:631–642.

51. Datsenko KA, Wanner BL. 2000. One-step inactivation of chromosomal genes in Escherichia coli K-12 using PCR products. Proc Natl Acad Sci 97:6640–6645.

52. Hall BG, Acar H, Nandipati A, Barlow M. 2014. Growth Rates Made Easy. Mol Biol Evol 31:232–238.

53. Okoye K, Hosseini S. 2024. R Programming, Statistical Data Analysis in Research 279– 303.

